# Flow cytometry method for absolute counting and single-cell phenotyping of mycobacteria

**DOI:** 10.1101/2021.05.01.442251

**Authors:** David A. Barr, Charles Omollo, Mandy Mason, Anastasia Koch, Robert J. Wilkinson, David G. Lalloo, Graeme Meintjes, Valerie Mizrahi, Digby F. Warner, Gerry Davies

**Affiliations:** Wellcome Centre for Infectious Diseases Research in Africa (CIDRI-Africa), Institute of Infectious Disease and Molecular Medicine, University of Cape Town, Observatory 7925, Cape Town, South Africa; SAMRC/NHLS/UCT Molecular Mycobacteriology Research Unit, DST/NRF Centre of Excellence for Biomedical TB Research, Institute of Infectious Disease and Molecular Medicine, Division of Medical Microbiology, Department of Pathology, University of Cape Town, Cape Town, South Africa; Institute of Infection and Global Health, University of Liverpool, Liverpool L7 3EA, UK; Liverpool School of Tropical Medicine, Pembroke Place, Liverpool L3 5QA, UK; Department of Medicine, University of Cape Town, Cape Town, South Africa; The Francis Crick Institute, London NW11AT, UK; Department of Medicine, Imperial College, London, W12 0NN, United Kingdom

## Abstract

Detection and accurate quantitation of viable *Mycobacterium tuberculosis* is fundamental to understanding mycobacterial pathogenicity, tuberculosis (TB) disease progression and outcomes; TB transmission; drug action, efficacy and drug resistance. Despite this importance, methods for determining numbers of viable bacilli are limited in accuracy and precision owing to inherent characteristics of mycobacterial cell biology – including the tendency to clump, and “differential” culturability – and technical challenges consequent on handling an infectious pathogen under biosafe conditions. We developed an absolute counting method for mycobacteria in liquid cultures using a bench-top flow cytometer, and the low-cost fluorescent dyes Calcein-AM (CA) and SYBR-gold (SG). During exponential growth CA+ cell counts are highly correlated with CFU counts and can be used as a real-time alternative to simplify the accurate standardisation of inocula for experiments. In contrast to CFU counting, this method can detect and enumerate cell aggregates in samples, which we show are a potential source of variance and bias when using established methods. We show that CFUs comprise a sub-population of intact, metabolically active mycobacterial cells in liquid cultures, with CFU-proportion varying by growth conditions. A pharmacodynamic application of the flow cytometry method, exploring kinetics of fluorescent probe defined subpopulations compared to CFU is demonstrated. Flow cytometry derived *Mycobacterium bovis BCG* time-kill curves differ for rifampicin and kanamycin versus isoniazid and ethambutol, as do the relative dynamics of discrete morphologically-distinct subpopulations of bacilli revealed by this high-throughput single-cell technique.

## Introduction

For more than 100 years, counting Colony Forming Units (CFU) has been the gold-standard for quantifying viable *Mycobacterium tuberculosis (Mtb)* bacilli, both *in vitro* and *ex vivo.* However, despite being a methodological foundation underpinning our scientific knowledge of *Mtb*, CFU counting has several technical and practical limitations, including cost and biosafety implications of maintaining multiple secondary cultures, time interval to results, loss of results from contamination, the inability to distinguish single cells from cell aggregates (clumps), and high intra- and inter-laboratory variation.^1–3^

More fundamentally, under some conditions sub-populations of viable *Mtb* cells do not form colonies and are therefore unobserved by CFU counting.^4–6^ This is of particular relevance to tuberculosis (TB) diagnostics and research because of the prevailing theory that the existence of phenotypically heterogenous sub-populations of bacilli – with differential metabolic or growth states – underlie profound aspects of TB disease biology, such as latency and the need for prolonged therapy to effect sterilising cure.^7–10^

Improved methods for absolute counting of mycobacteria and phenotypic characterisation of subpopulations are therefore desirable. Flow cytometry (FCM) is a well-established technique for counting and characterising eukaryotic cells, and its potential to advance single-cell analyses in microbiology has been discussed in depth.^11,12^ Several groups have applied FCM to mycobacteria, including drug sensitivity testing,^13–22^ investigation of cell biology, ^23–28^ early phase diagnostic test development,^29,30^ live/dead discrimination,^31,32^ and more advanced single-cell phenotyping.^23,33,34^ Fluorescent dyes used by prior investigators include: probes of membrane integrity (the nucleic acid stains SYTOX-green,^33^ SYTO-9,^24,31,32^ SYTO-BC,^24,32^ SYTO-16,^16,28^ SYBR-green I,^30^ propidium iodide,^16,24,31,32,35^ TO-PRO-3 iodide,^34^ and auramine-O^13,35,36^); probes of metabolic activity (esterase substrate dyes fluorescein diacetate,^14,17,20,21,37^ and Calcein-violet^33^) and membrane potential (diethyloxacarbocyanine iodide ^25,34^ & rhodamine-123^27^). In general, absolute bacillary counts have not been derived from FCM; instead, batch measures of fluorescence *(e.g.* mean fluorescence signal), ^13–17,20,21,27,29,37^ qualitative read-outs (*e.g.* scatter-plots),^18,26,34^ or percentages^23,25,33^ are reported. Growth conditions, processing (*e.g.* washes and fixation), and staining protocols vary widely. In the few cases in which the same stains have been used by different groups, results are often contradictory: for example, propidium iodide is reported to stain 0% of heat killed *M. tuberculosis* by one study,^38^ and 100% by another.^31^ No comparisons of FCM counts with CFU enumerations have been published.

The aims of the current study were:

1. To develop and validate a method for absolute counting of mycobacteria *in vitro* using FCM.
2. To explore the use of fluorescent dyes as probes of cell function to define subpopulations of bacilli in discrete physiological states.
3. To compare dynamics of FCM-defined subpopulations and CFU in liquid cultures over time (growth curves), and over time in the presence of antimycobacterial compounds (time-kill curves).

We report a method developed on a low-cost flow cytometer (BD Accuri™ C6), using two commercially available fluorescent dyes (SYBR®-Gold and Calcein-AM). The BD Accuri C6 flow cytometer has fixed alignment and pre-optimised detector settings, can record volume of sample processed without use of counting beads, and is small enough to fit on a benchtop or inside a bio-containment hood. SYBR®-Gold (SG), a proprietary cyanine dye (excitation ~495nm, emission ~573nm) with >1000-fold fluorescence enhancement when bound to nucleic acid, was designed for use in gel electrophoresis.^39^ SG has previously been shown to have substantially greater sensitivity than auramine-O for quantitative fluorescence microscopy of heat-fixed mycobacteria (99% versus 65-80%),^40^ but it has not been applied in FCM. Calcein-AM (CA) is a non-polar, lipophilic ester which becomes charged and fluorescent when hydrolysed by ‘house-keeping’ esterases ubiquitous in the cytoplasm of living cells.^41^ Hendon-Dunn and colleagues previously showed that the fluorescence of mycobacteria stained with Calcein-violet-AM correlated with rate of growth in a chemostat and declined with antimicrobial killing of bacilli.^33^

In the present study, we applied SG staining after heat killing bacilli to define a total intact cell count denominator; SG staining without heat killing to probe cell membrane integrity as a marker of death or damage; and CA staining without heat killing to probe metabolic activity as a marker of vitality.

## Results

### Setting fluorescence threshold values for FCM events improves validity of absolute bacilli counts

The BD Accuri C6 flow cytometer has a fixed dynamic range for voltage and gain, but allows thresholds to be set on two signal values from light scatter and/or fluorescence channels. Signals below the set threshold are not recorded as events. For absolute counting of cells, an optimal threshold is one that is not so high as to exclude true events (signals from cells), yet high enough that it excludes electronic noise and signal from debris (which can otherwise mask true events owing to the refractory period of photodetectors). Typically, thresholds are set on forward and/or side scatter of light (FSC and SSC), as this allows fluorescence-positive and -negative events to be recorded without biasing measurements from the fluorescence channels.

We investigated different threshold strategies as follows. Mid-log phase *M. bovis* BCG cultures were analysed after 2-fold dilution in 0.15% v/v Tween80 PBS solution. These were compared to an identical preparation of cell-free 7H9 broth as a negative control. Permutations of threshold settings were screened. In each case a gate was set around an apparent discrete population of events visible on a log(SSC) by log(FSC) plot, with the gate set manually to minimise the ratio between negative control and the paired BCG sample event counts. The optimal ratio (false positive event count in cell free broth divided by the paired BCG culture count) was defined as the false discovery rate; therefore, threshold parameters which maximised the absolute count in the BCG broth gate and minimised the false discovery rate were sought.

Optimal threshold values based on light scatter (FSC and SSC) were inconsistent across replicates and were never associated with false discovery rates less than 10%. By contrast, thresholding on SSC and fluorescence (FL1 533/30 nm) in heat-killed, SYBR-gold stained mid-log BCG consistently reduced the false discovery rate to <0.5%, at the same time increasing absolute cell counts by more than one logarithm compared with thresholding on light scatter alone on the same samples (figure 1). Further, this strategy reduced the coefficient of variation between technical replicates to <5%, and gave near perfect linearity across serial dilutions of the same sample (R^2^>0.99) (figure 1).

**Figure 1.**
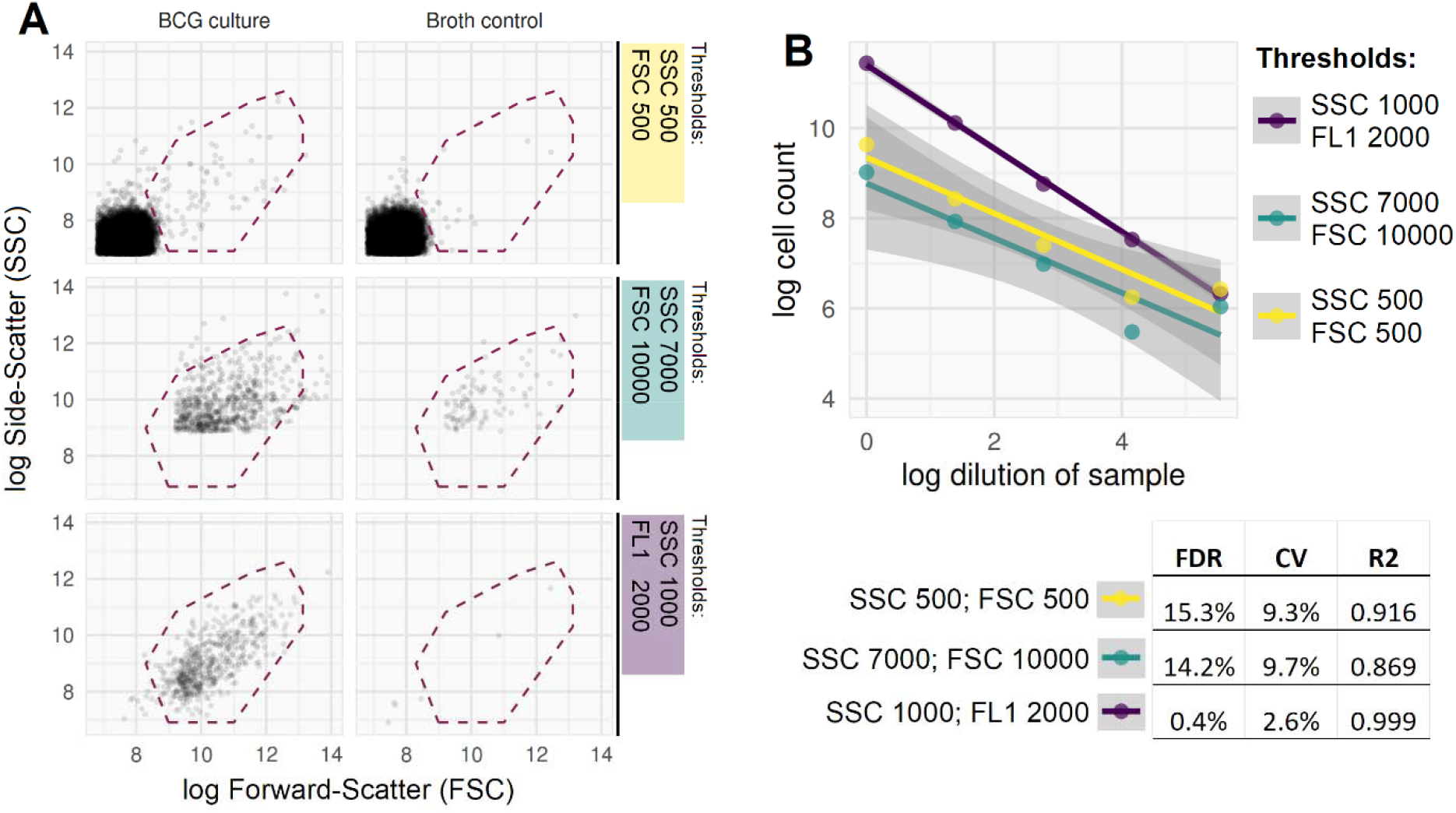
Comparison of 3 thresholding strategies in heat killed and SYBR-gold stained mid-log phase BCG broth culture. **A**. Example FCM plots for three thresholding strategies (rows 1-3) applied to *M. bovis* BCG broth culture (column1) and cell-free broth (cell-free negative control, column 2). Counts are extracted for the gated population (events within dashed red line), which is placed to maximise the count in BCG broth and minimise the count in the cell-free control. Recorded events in the low light scatter value thresholding (first row) are dominated by debris/noise, seen as a dense population with low SSC and FSC values in lower left quadrant; this is equally apparent in the cell-free control. Higher light scatter thresholding (second row) excludes these events, but still records a substantial portion of higher SSC/FSC noise (seen in cell-free control), and the threshold level appears to bisect the ‘real’ cell population; *i.e.*, losing cells from analysis. By contrast, the thresholding based on fluorescence (third row) is qualitatively better, with very few false positive events in the cell-free control, and detection of a discrete cell population in BCG broth which is not artificially bisected. **B**. Greater internal consistency in the FL1/SSC thresholding strategy, with less error across serial dilutions of a *M. bovis* BCG culture. Quantitative evidence of improved absolute count validity includes a lower false discovery rate (FDR, defined as false positive cell count in cell-free control divided by paired cell count from broth); lower coefficient of variation (CV, calculated by standard deviation/mean from 5 technical replicates, averaged for 3 biological replicates); and higher R^2^ from linear fit across serial dilution series (one biological replicate as shown in figure; p<0.001 for F test comparing FL1/SCC to either FSC/SSC strategy; 95% confidence intervals for linear fit shown with grey shaded areas).

### Clumping in mycobacterial broth cultures can be observed and quantified using FCM

In all mycobacterial broth cultures tested – *M. bovis* BCG, *M. tuberculosis*, and *M. smegmatis*, a second population of events with higher FSC and SSC became evident from early log-phase onwards, developing into the dominant population in mid- or late log-phase (figure 2A). To investigate the nature of these distinct populations, events gated on the two light-scatter populations were sorted for downstream microscopy (figure 2 B&C). This analysis revealed that the higher light-scatter population was composed of clumped cells, despite the fact that all cultures were grown in detergent (Tween80, 0.1% to 0.25% v/v) under continuous agitation (150 to 200 rpm), and notwithstanding the use of sonication prior to flow cytometry.

**Figure 2.**
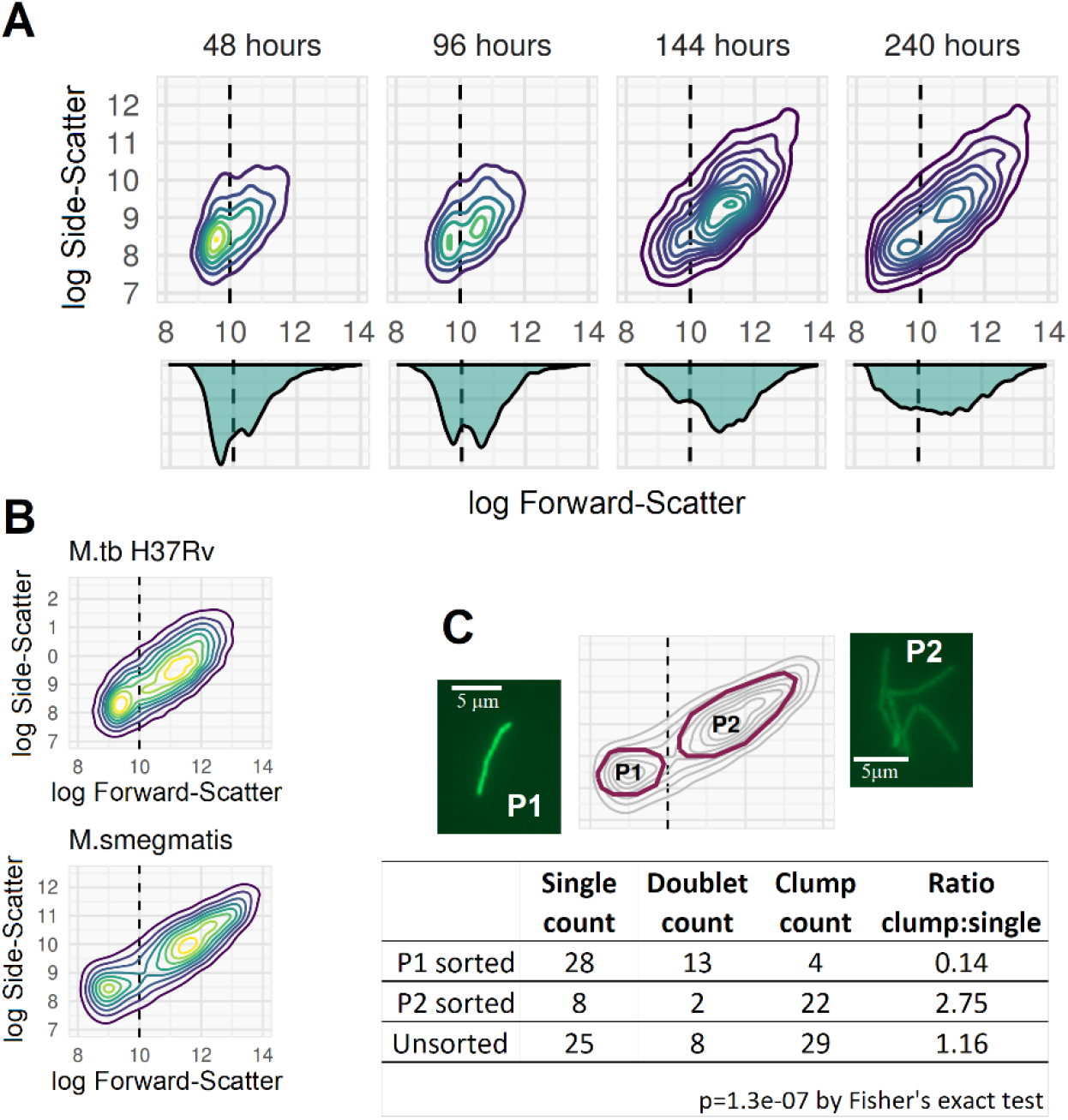
Identifying mycobacterial clumping with FCM. **A**. 2D-density plots of FSC v SSC on log scale for a culture of *M. bovis* BCG grown with 0.15% v/v Tween80 at 180 rpm. Plots are from samples taken at 48, 96, 144, and 240 hours after bacilli were sub-cultured from a log-phase starter culture into pre-warmed broth (early, early-mid, late-mid and late log-phase, respectively). Samples were diluted 10-fold or 100-fold (later samples) in 0.25% v/v Tween80 PBS and sonicated for 60 seconds prior to flow cytometry. The tail of higher SSC and FSC events at 48-hours is seen to develop into a discrete second subpopulation by 96-hours, which continues to expand into a higher SSC and FSC region and become the predominant subpopulation by the end of log-phase. All plots are constructed from 5000 events. **B**. *M. tuberculosis* and *M. smegmatis* processed as above (both mid-late log phase) also develop dual populations separating on FSC and SSC, replicating the *M. bovis* BCG findings. **C**. *M. smegmatis* sample was run on a BioRad S3 cell sorter with P1 and P2 sorted for downstream fluorescence microscopy (representative images shown). *M. smegmatis* was used for cell sorting owing to concerns about aerosolising *M. tuberculosis* or *M. bovis* BCG. P1 comprised majority single cells or doublets, while P2 comprised majority clumps (manually quantified from fluorescence microscopy images).

### Clumping is a major determinant of CFU count& can be controlled by needle emulsification, but not vortex, sonication or centrifugation

Having established the ability to quantify mycobacterial clumping in broth cultures using FCM, we next tested the comparative efficacies of standard microbiological methods for clump dispersal. Vortex and sonication failed to disrupt the clumped population observed on FCM; by contrast, needle emulsification of the broth culture largely eliminated clumping (figure 3A). Disruption of clumps by needle emulsification increased the single-cell population seen on FCM, and therefore the CFU count, by more than 0.5 log (figure 3B). Larger clumps not disrupted even by needle-emulsification emerged in late-stage broth cultures (figure 4).

**Figure 3.**
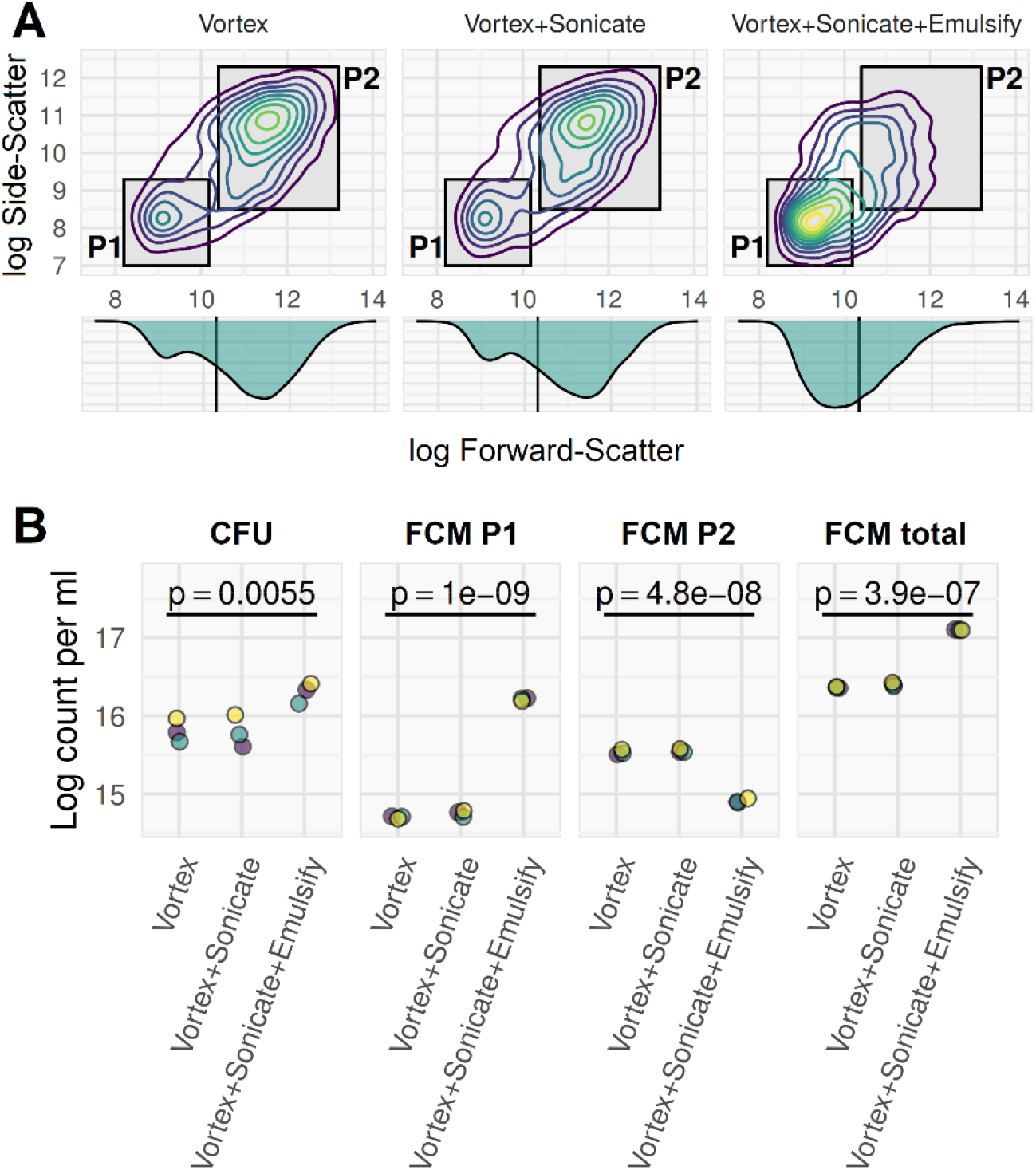
Needle-emulsification, but not vortex or sonication, disrupt clumps and increases cell counts. Mid-log phase culture of *M. smegmatis* grown in 0.15% v/v Tween80 7H9 with 150 rpm agitation and diluted 10-fold in 0.15% v/v Tween80 PBS and stained with Calcein-AM prior to FCM on a BD Accuri C6. Data acquisition with thresholds SSC>1000 and FL1>2000. Samples were processed by 60 second vortex, or by 60 second vortex followed by 5 minutes sonication in water bath, or by both these methods followed by needle emulsification (**12** passes through a double Luer lock-ended, 25 Gauge, 4-inch, micro-emulsifying needle with a reinforcing bar (Cadence Inc.). **A.** Two populations are seen which are differentiated by light-scatter: single cells (P1) and clumps (P2). Qualitatively, vortex and sonication processing did not disrupt P2 population (clumps), but needle emulsification (far right plot) shifted events from predominantly P2 (clumps) to predominantly P1 (single cells). **B**. Counts of CFUs or FCM events with three-replicates from 3 independent cultures (purple, green, yellow dots). Emulsification resulted in a greater number of CFU and decreased the P2 count while increasing the P1 count substantially. The apparent total cell counts were increased by emulsification by an order of magnitude: both CFU count and total flow cytometry CA positive count increased by half to one unit on log scale. This can be interpreted as resulting from clumps (P2 population) being disaggregated into single cells (P1). The p-values were determined from repeated-measures ANOVA by cell-disruption method.

**Figure 4.**
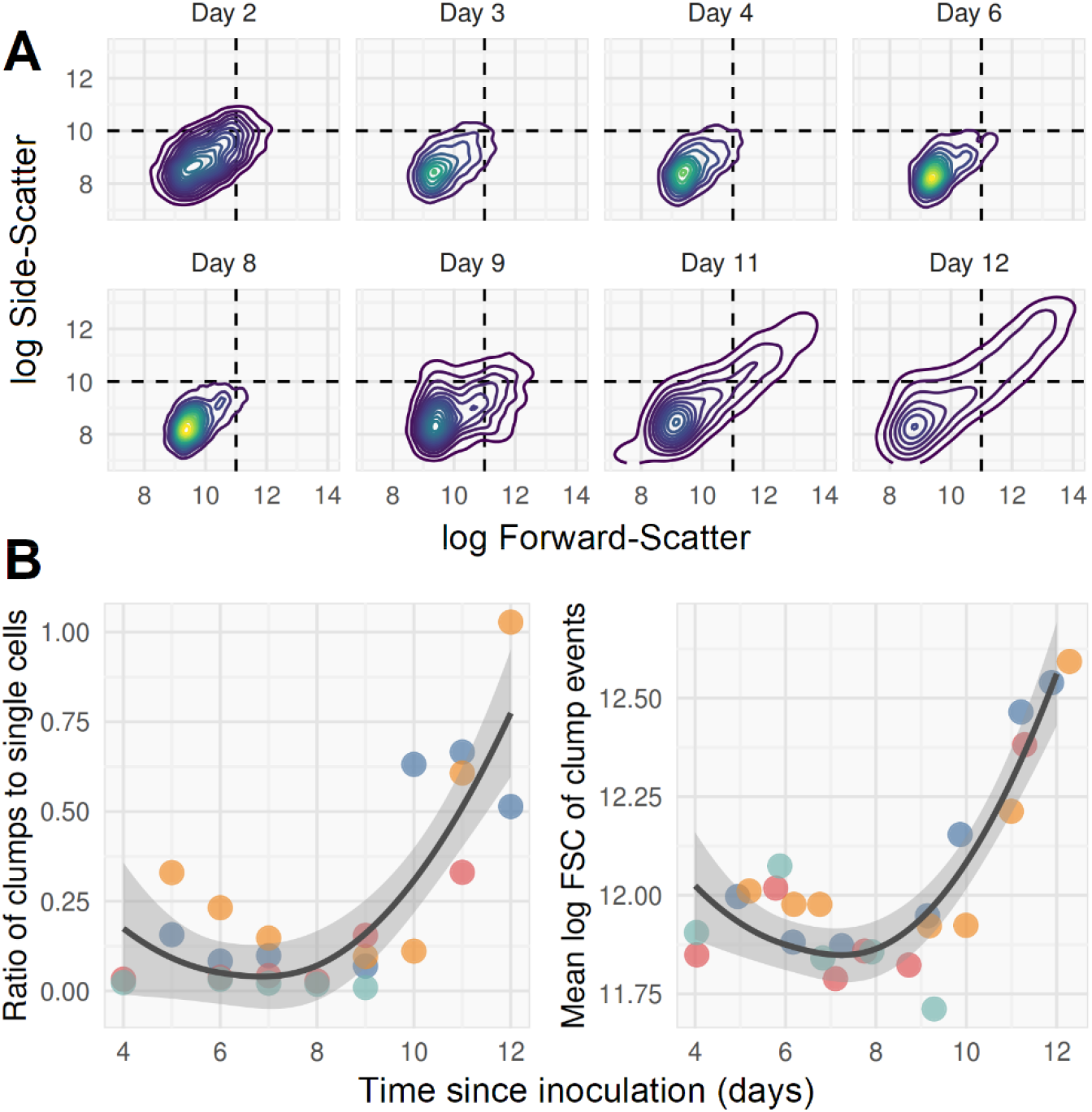
Clumps which are resistant to disruption by emulsification eventually emerge in late log-phase cultures. **A**. *M. bovis* BCG culture in 0.15% v/v Tween80 7H9 with 150 rpm agitation and diluted 10-fold in 0.15% v/v Tween80 PBS before bacilli were heat-killed and stained with SYBR-gold. Samples were needle-emulsified (**12** passes through a double Luer lock-ended, 25 Gauge, 4-inch, micro-emulsifying needle) prior to FCM on BD Accuri C6; data acquisition with thresholds SSC>1000 and FL1>1000. Timepoints are days post inoculation into pre-warmed broth from log phase starter culture. A long tail of clumps, extending into the upper-right quadrant of higher SSC and FSC, emerges from around day 9, at OD_600_ ~ 0.3. Clumps were defined as events with SSC and FSC values greater than 10-log and 11-log (events in upper right quadrant of plots). **B**. Clumps and single cells quantified by flow cytometry (four independent replicates of data represented by A; replicates are shown with different colours). Emulsification appears able to disrupt clumps until late log phase (~day 8), when both the ratio of clumps to single cells and size (approximated by mean FSC) of clumps rise rapidly. Line of best fit with 95% confidence interval band is a LOESS regression line ignoring dependence by replicate.

A standard method for preparation of single-cell suspensions of mycobacteria is centrifugation, based on the premise that cell clumps are selectively pelleted by gravity, with single cells remaining in suspension. ^42^ However, using FCM we found that the ratio of clumps to single-cells was unaffected by centrifugation (figure 5).

**Figure 5.**
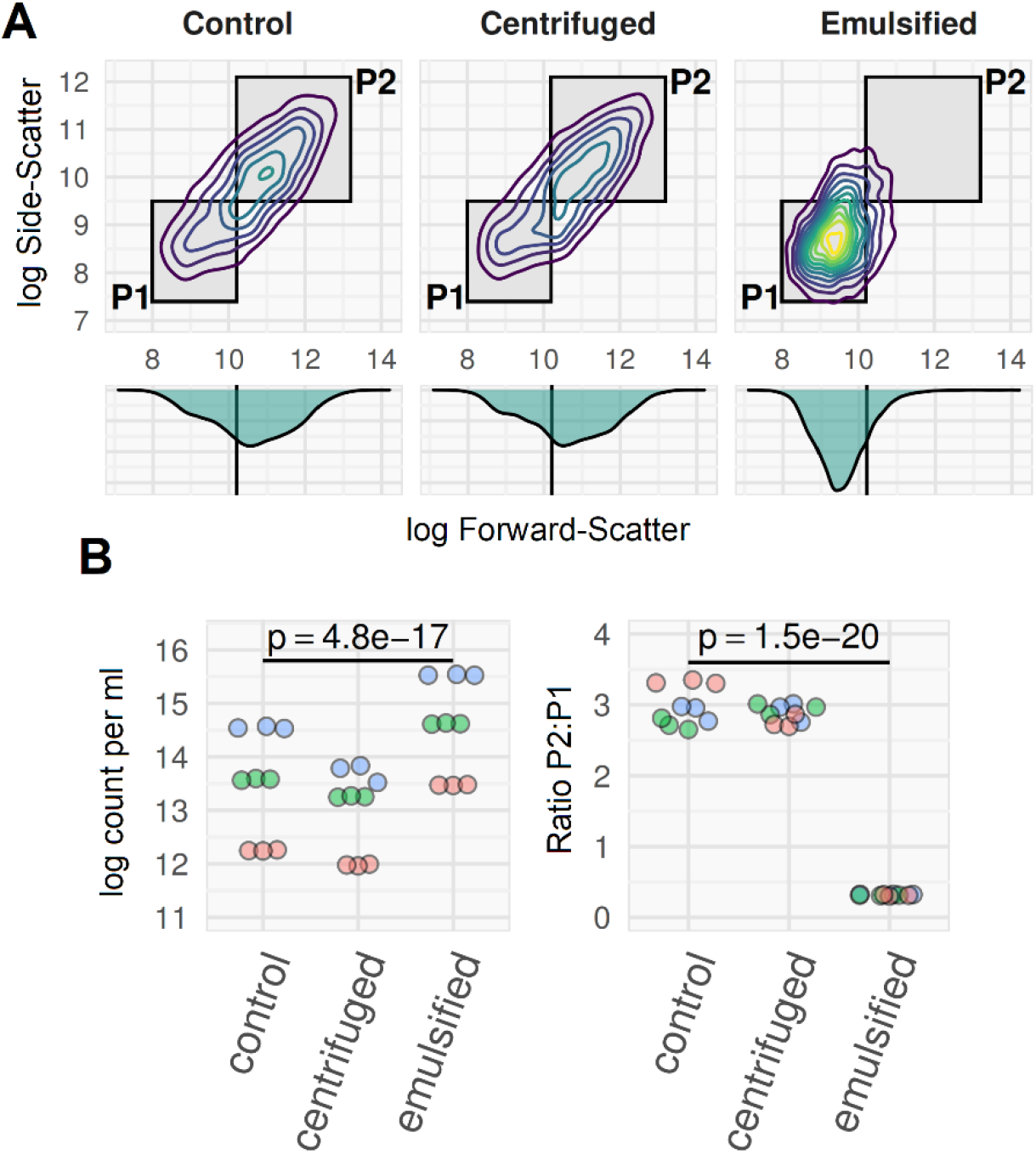
Ratio of clumped to single-cell *M. smegmatis* is not altered by low-*g* centrifugation. Three mid-log phase *M. smegmatis* cultures grown in 0.05% Tween80 7H9 with continuous agitation at 150 rpm(3 biological replicates shown in blue, green and red), processed three ways. **Control sample**: 10^−1^ dilution in 0.1% Tween80 PBS, no physical disruption. **Centrifuge sample:** 10ml + 5ml 0.1% Tween80 PBS; spun in 15ml centrifuge tubes at 120 x *g* for 8 minutes with no brake. **Emulsified sample**: 10^−1^ dilution in 0.1% Tween80 PBS, 12x needle emulsified. Supernatant used for counts, as per ref 42 main manuscript. All samples heat-killed and stained with SYBR-gold prior to flow cytometry on BD Accuri-C6, with thresholding on SSC and FL1. Data are for three technical replicates of each culture. **A**. Qualitatively, the FSC by SSC flow plots were similar for centrifuge method and control, compared to the needle-emulsified sample where the cell-clump population (P2) was not evident. **B**. The ratio of clumps to single cells (p2:p1) was the same in the control and centrifuge preparations, but was much lower (and with less variation across replicates) with emulsification. Apparent cell counts were lower with centrifugation (owing to loss of cells in pellet) and higher with emulsification (owing to disruption of clumps). Three independent culture replicates (read, blue, and green; each processed 3 times in each condition for technical replicates); p-values from repeated measures ANOVA (technical replicates nested within culture replicates).

### Growth dynamics of FCM-defined bacilli populations compared to CFU

A FCM protocol for absolute counting of bacilli – incorporating SYBR-gold or Calcein-AM staining, needle-emulsification to disperse clumps, and thresholding on fluorescence (summarised in figure 6) – was used to explore dynamics of *M. bovis*BCG growth in broth culture. We defined three FCM populations using this protocol:

1. **Calcein-AM-positive (CA+)**–live sample stained with Calcein-AM, to give an esterase positive, or ‘metabolically active’, cell count.
2. **SYBR-gold-positive (SG+)**–live sample stained with SYBR-gold, to count cells which have membranes permeable to SYBR-gold, implying membrane damage.
3. **Heat-killed ‘total cell count’ (HK)**–sample incubated in water-bath at 60°C for 12 minutes to permeabilise cell-membranes, followed by SYBR-gold staining. This is proposed to give a total count of intact cells containing nucleic acid, and therefore provides a denominator for calculating the proportion of cells which are CA+, or SG+, or colony-forming. The selected heat-kill time and temperature were selected as the minimum to reliably maximise the HK count.

**Figure 6.**
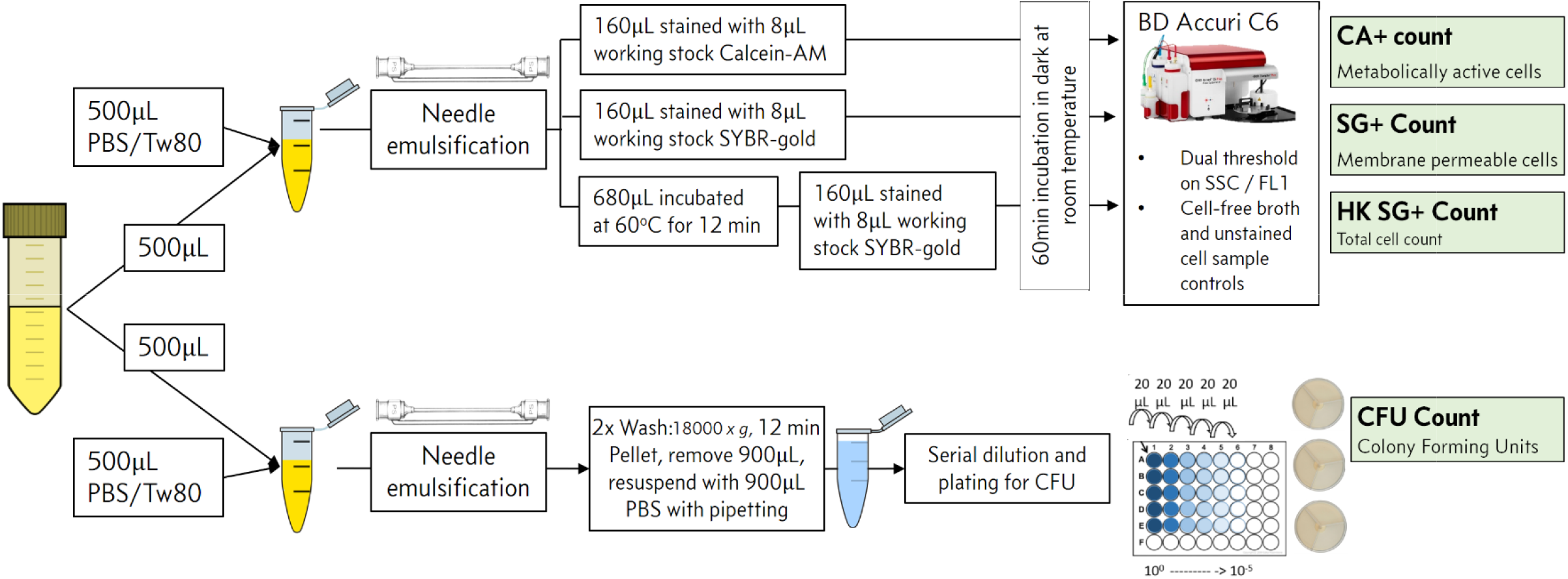
Schematic for final FCM count SOP used in *M. bovis* BCG culture growth & time-kill dynamics experiments

CFU counts after needle emulsification were determined in parallel (figure 6).

In all culture growth phases (lag, log, stationary), the HK cell count was greater than CA+, SG+, or CFU counts (figure 7A), and was accepted as a total cell count. During log-phase, most cells were CA+ and colony forming, with SG+ cells constituting a minority sub-population (figure 7B-D). When entering stationary phase, CA+ and CFU counts started to fall, with a simultaneous rise in SG+ cells, which subsequently became the dominant subpopulation (figure 7B-D).

**Figure 7.**
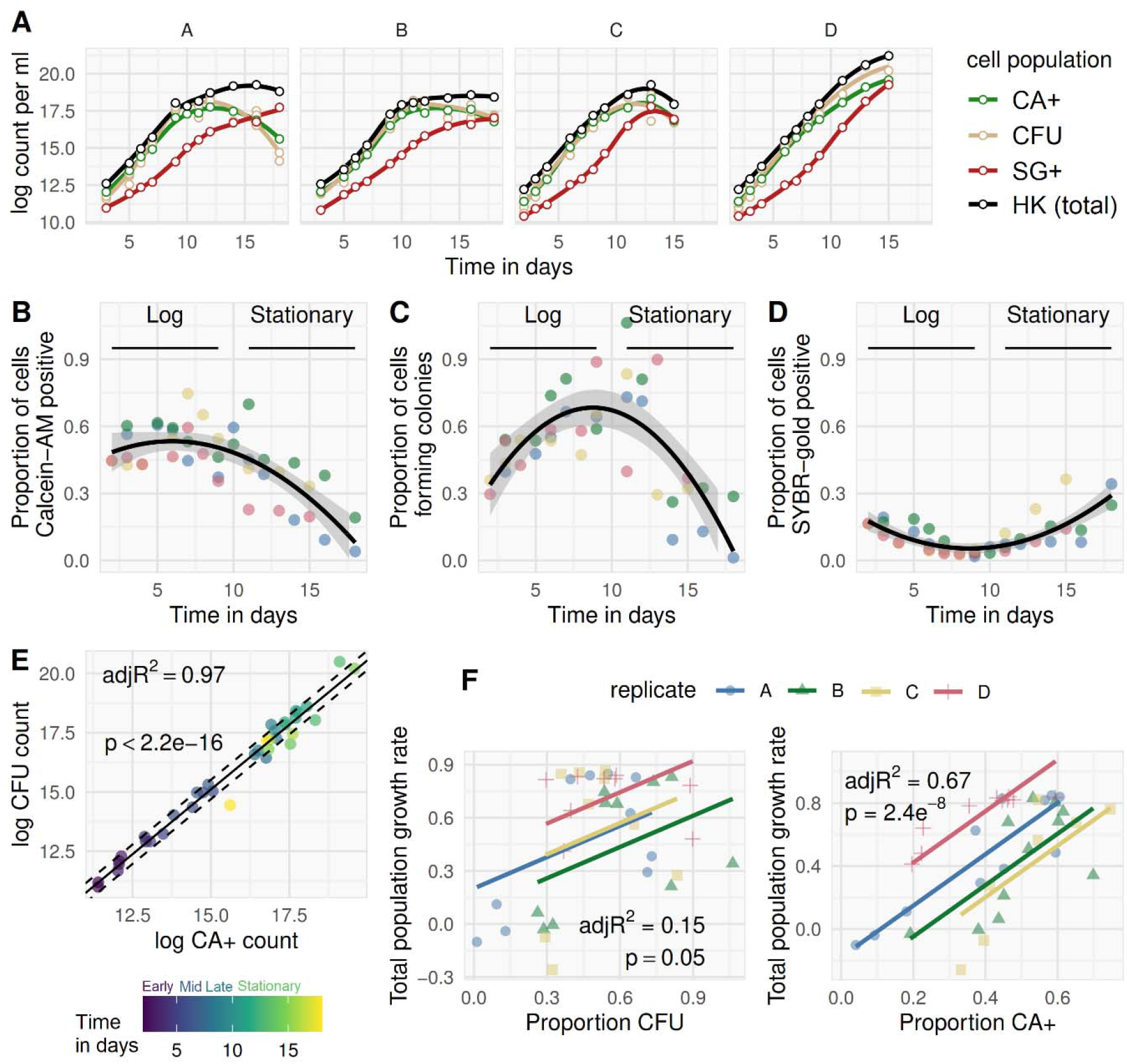
Bacillary population dynamics during standard growth in culture. Four independent *M. bovis* BCG cultures (replicates i to iv), grown in 50ml 0.15% v/v Tween80 7H9, 500ml tissueculture flasks at 150 rpm agitation, were serially quantified by CFU and FCM counts (method outlined in figure 6) between 2 and 15 days after inoculation into pre-warmed broth from a log phase starter culture. (**A**) Using the heat-killed SYBR-gold stained (HK) cell count as total cell denominator, the proportion of bacilli which were Calcein-AM positive (CA+), colony-forming (CFU), and permeable to SYBR-gold without heat-killing (SG+) are shown over time post inoculation (**B-D**); each replicate is plotted using a different colour; LOESS line-of-best-fit and 95% CI shown for the observations aggregated across replicates. Linear correlation between log CA+ and log CFU counts was strong (**E**), but with a dependency on phase of growth (time in days from inoculation shown by colour; non-constant variance (NCV) test for heteroscedasticity, p=0.03, dashed lines are +/- 1 SD of residual variation). Rate of population growth is defined as instantaneous rate of change in total cell count (slope of the tangent to the curve at a given timepoint; *i.e.*, first derivative of the growth curve). Rate of total population growth was regressed on proportion of bacilli able to form colonies, or on proportion CA+ (**F**), at any given timepoint, with each replicate (i to iv, again shown by colour) allowed to differ by intercept but not slope.

Correlation between CFU and CA+ counts was growth-phase dependent (figure 7E) with close co-variance in early-to-mid-log phase progressively diminishing in late-log (when CFU > CA+) and stationary phase (when CA+ > CFU). ‘Total population growth rate – defined using the instantaneous rate-of-change of the log HK total cell count (the slope of the tangent to the curve at a given timepoint, *i.e.* the first-derivative) – correlated with the proportion of bacilli which were CA+, but not the proportion of bacilli forming colonies (figure 7F).

### *In vitro* pharmacodynamics of *M. bovis* BCG by CFU and FCM counting

Having established growth dynamics in the absence of antimicrobials, the FCM count method was applied to pharmacodynamic (PD) time-kill analysis of *M. bovis* BCG. Starter cultures (100 ml in 500ml tissue culture flasks containing 0.15% v/v Tween 80 7H9 medium) were grown to a density of ~2×10^5^ CA+ cells per ml, then split into 20ml samples in 50ml conical flasks. Antimicrobials (rifampicin, isoniazid, kanamycin, ethambutol) were added at a range of final concentrations in multiples of their minimum inhibitory concentration (MIC99), and bacilli quantified at 0, 24, 48, 72 and 120 hours using FCM and CFU counting. The experiment was repeated on three separate occasions to ensure independent biological replicates.

Raw FCM data plots for heat-killed, Calcein-AM, and SYBR-gold stained preparations from one of three independent replicates are shown for selected conditions in figure 8A-C. Time-kill curves based on absolute counts, and proportions (CA+/HK, SG+/HK, CFU/HK counts), are shown in figures 9A&9B, respectively.

**Figure 8.**
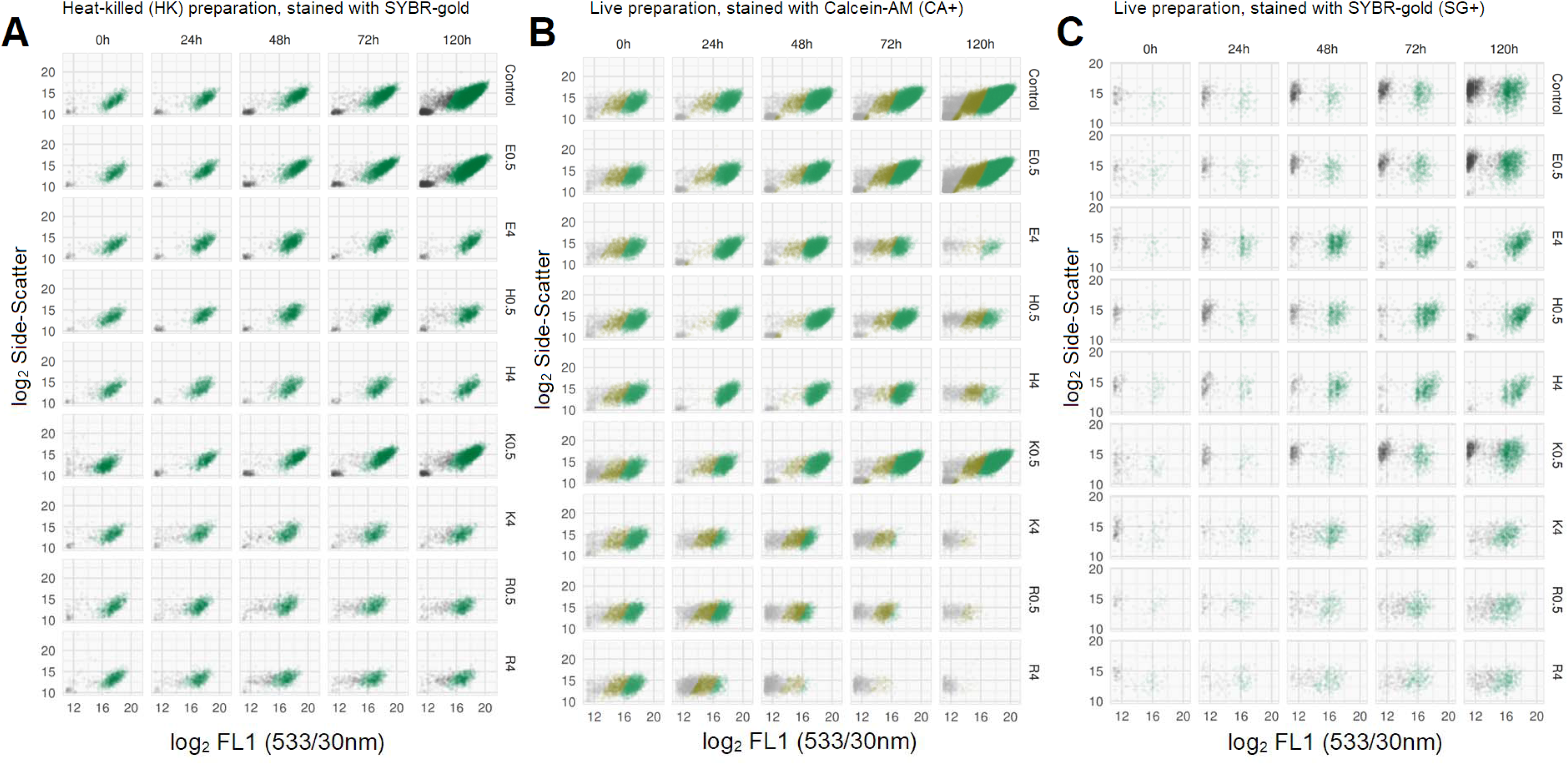
Raw FMC plots of HK, CA+, and SG+ events for selected antimicrobial conditions and timepoints. Raw FMC data from one of three independent replicates. SYBR-gold stained heat-killed (HK) samples (A, left), Calcein-AM stained live samples (B, middle), and SYBR-gold stained live samples (C, right), by antimicrobial condition (rows) and time-point (hours) post introduction of antimicrobials (columns). Antimicrobial indicated by letter prefix (E, ethambutol; H, isoniazid; K, kanamycin; R, rifampicin) and concentration in multiples of MIC_99_ by suffix letter (*e.g*., R4 = rifampicin at 4x MIC). FCM events in each plot are coloured by K-means clustering on all light-scatter and fluorescence dimensions – an unsupervised classification (machine learning) algorithm used to define subpopulations without subjective manual placement of gates.

**Figure 9.**
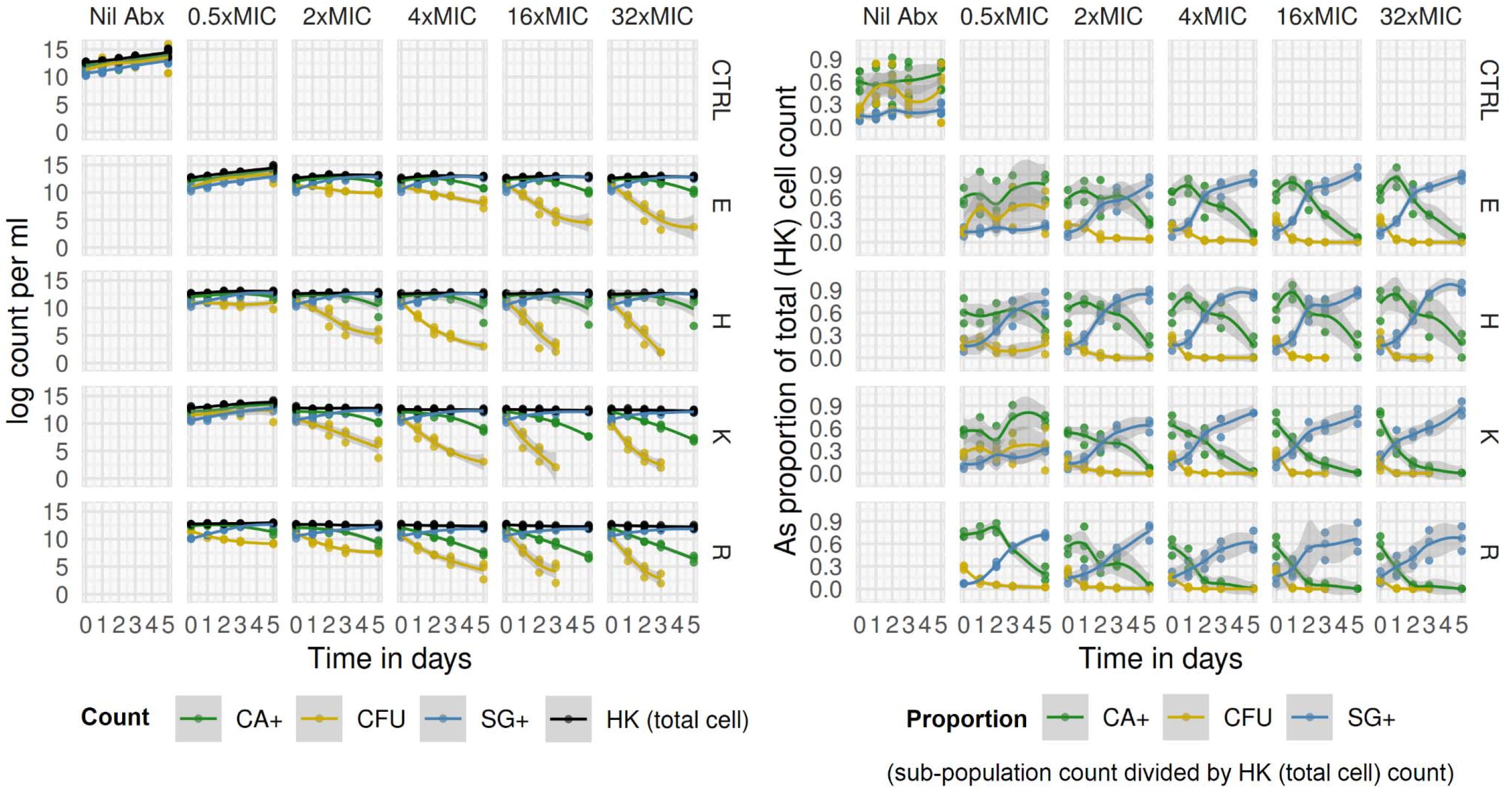
Time-kill curves generated through FCM defined cell populations and CFU counts. Data from three replicates. Counts (left panel) and proportions (right panel) by cell population [CA+ = Calcein-AM-stained in live samples; CFU = colony forming units; SG+ = SYBR-gold-stained in live samples; HK = SYBR-gold-stained in heat-killed samples] over time by antimicrobial condition. Antimicrobial indicated by letter in rows (E, ethambutol; H, isoniazid; K, kanamycin; R, rifampicin; CTRL, antimicrobial-free [“Nil Abx”] broth) and concentration in multiples of MIC in columns. Non-parametric Loess regression line and shaded 95% confidence intervals shown. Proportions are derived using HK count as a total cell count denominator *(i.e.* CA+/HK, SG+/HK, CFU/HK).

Compared to antimicrobial-free controls, total cell count (HK count) growth was generally impeded by the presence of antimicrobials, although exponential growth still occurred with ethambutol and kanamycin at 0.5xMIC concentration of both antimicrobials. Critically, even at high concentrations of all antimicrobials tested, HK count did not show dramatic reduction over 120 hours of exposure.

By contrast, CA+ and CFU counts fell substantially over that time period. Notably, CFU counts declined earlier than CA+ counts for all antimicrobial tested for all inhibitory concentrations. Rifampicin or kanamycin exposure resulted in an earlier decline in CA+ counts than was observed for isoniazid or ethambutol, which at high concentrations showed an initial increase in proportion of cells CA+ at 24 hours, before a sustained fall to day 5.

Because a fraction of cells were SG+ under any condition, the major driver of absolute SG+ count was the total cell count, this can be seen in the antimicrobial-free controls where the highest SG+ counts were seen at late stages of growth. The proportion SG+ was, however, antimicrobial dependent: SG+ cells were a majority by day 5 in all supra-MIC concentration conditions, but the rise in the SG+ proportion occurred earlier and was larger for isoniazid and ethambutol than for rifampicin or kanamycin.

To summarise differing effects by antimicrobial and subpopulation, sigmoidal E_max_ models were fitted to the time-kill data (figure 10). Based on CFU time-kill curves, rifampicin, kanamycin, and isoniazid all have similar Emax values, while ethambutol is substantially lower. CA+ time-kill E_max_ was higher for rifampicin and kanamycin; and lower for ethambutol and isoniazid. The pattern was reversed for the effect on SG+ proportion. Finally, while the effects of antimicrobials on total cell count (HK count) were modest, they did differ by antimicrobial, with E_max_ highest for rifampicin and lowest for ethambutol.

**Figure 10.**
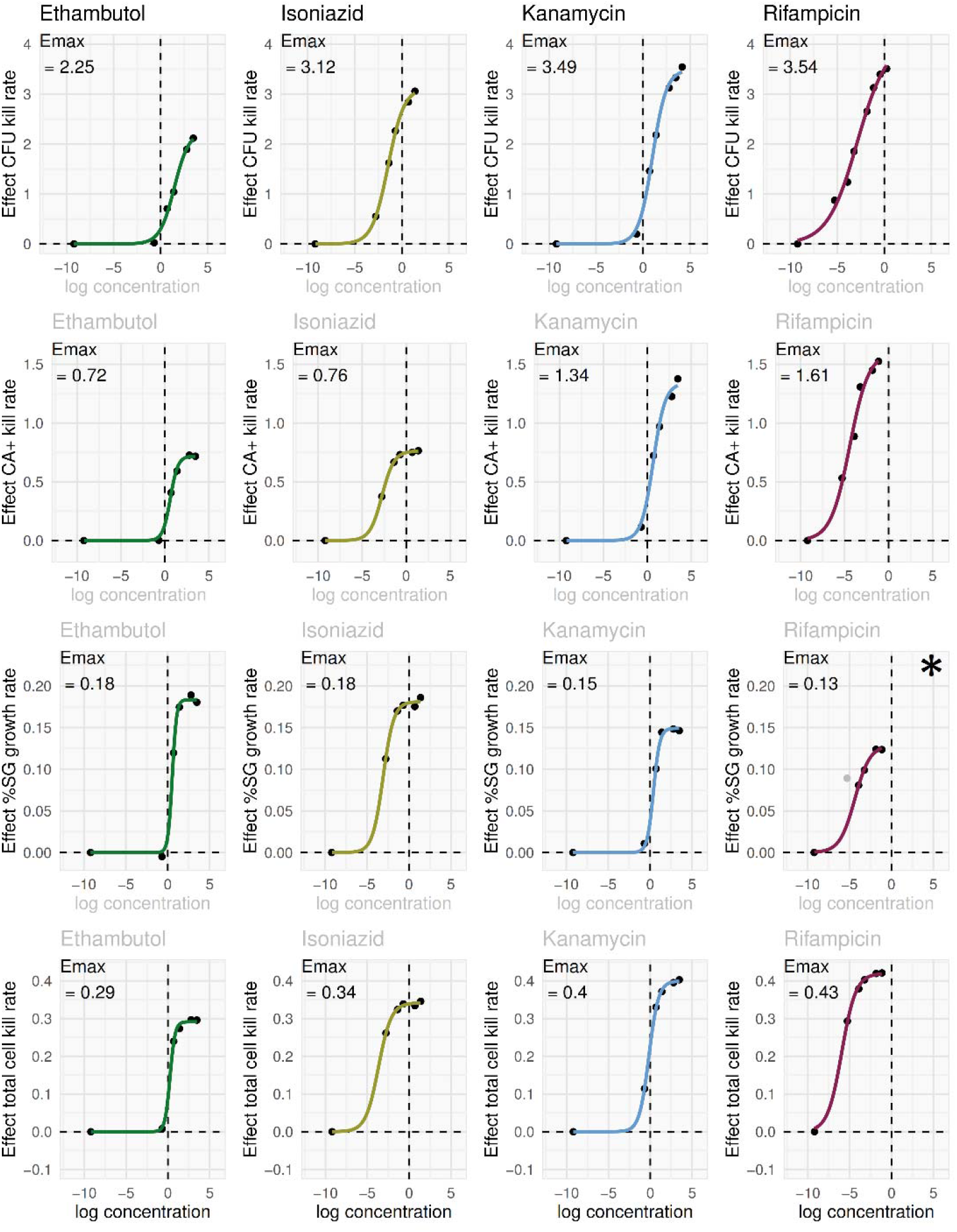
E_max_ models applied to time-kill data from FCM defined cell population and CFU counts. The time-kill data in figure 9 were modelled using a linear mixed-effects model to extract an estimate of monoexponential elimination rate for each antimicrobial condition as a summary pharmacodynamic (PD) measure for each FCM defined cell population and CFU counts. Antimicrobial effects extracted from these models were related to drug concentration for each antimicrobial using a standard sigmoid E_max_ PK/PD model, shown here for each antimicrobial (columns) and each cell population (rows). * A sigmoidal E_max_ model could not be fit for rifampicin %SG+ growth rate data due to non-convergence; the fit shown is from a model excluding the outlier data point at concentration 0.005 mg/ml (−5.3 on log scale). When this data point (indicated in grey) was excluded, the model converged.

### Subpopulations of cells by SYBR-gold staining characteristics

In addition to count data, qualitative differences in fluorescence were seen in live bacilli stained with SYBR-gold, with two subpopulations of SG+ cells separated by FL1 intensity (most distinct after 72 hours of isoniazid or ethambutol exposure, figure 8C). We hypothesised that two populations of bacilli with different SG staining properties were revealed by the membrane permeabilising effects of these antimicrobials. To investigate this possibility, we developed a protocol for permeabilising *M. bovis* BCG membranes without bacillary destruction (detailed in methods), and characterised these subpopulations under different antimicrobial conditions by quantifying them and through direct microscopy after cell-sorting.

Dual SG+ subpopulations were discriminated by distribution peaks (figure 11A) and were seen under all conditions, including growth without antimicrobial exposure (figure 11B) with one population (labelled P2) returning a mean fluorescence twofold higher than the other (labelled P1) (figure 11A&C). Fluorescent microscopy of cell-sorted samples showed that, compared to P1, P2 bacilli were longer (mean 4.0μm versus 2.5μm), with double the number of fluorescent foci (mean 6.1 versus 3.2) (figure 11D). The ratio of P2 to P1 cells in antimicrobial-free cultures was median 1.75, and non-significantly higher when bacilli were exposed to rifampicin or kanamycin (median 1.83 and 1.86, respectively), but significantly higher after exposure to ethambutol or isoniazid (median 2.11 and 2.00, respectively; p<0.001 for both by rank-sum test). The P2:P1 ratio when bacilli were incubated with both rifampicin and isoniazid matched rifampicin mono-exposure rather than isoniazid mono-exposure (figure 11E).

**Figure 11.**
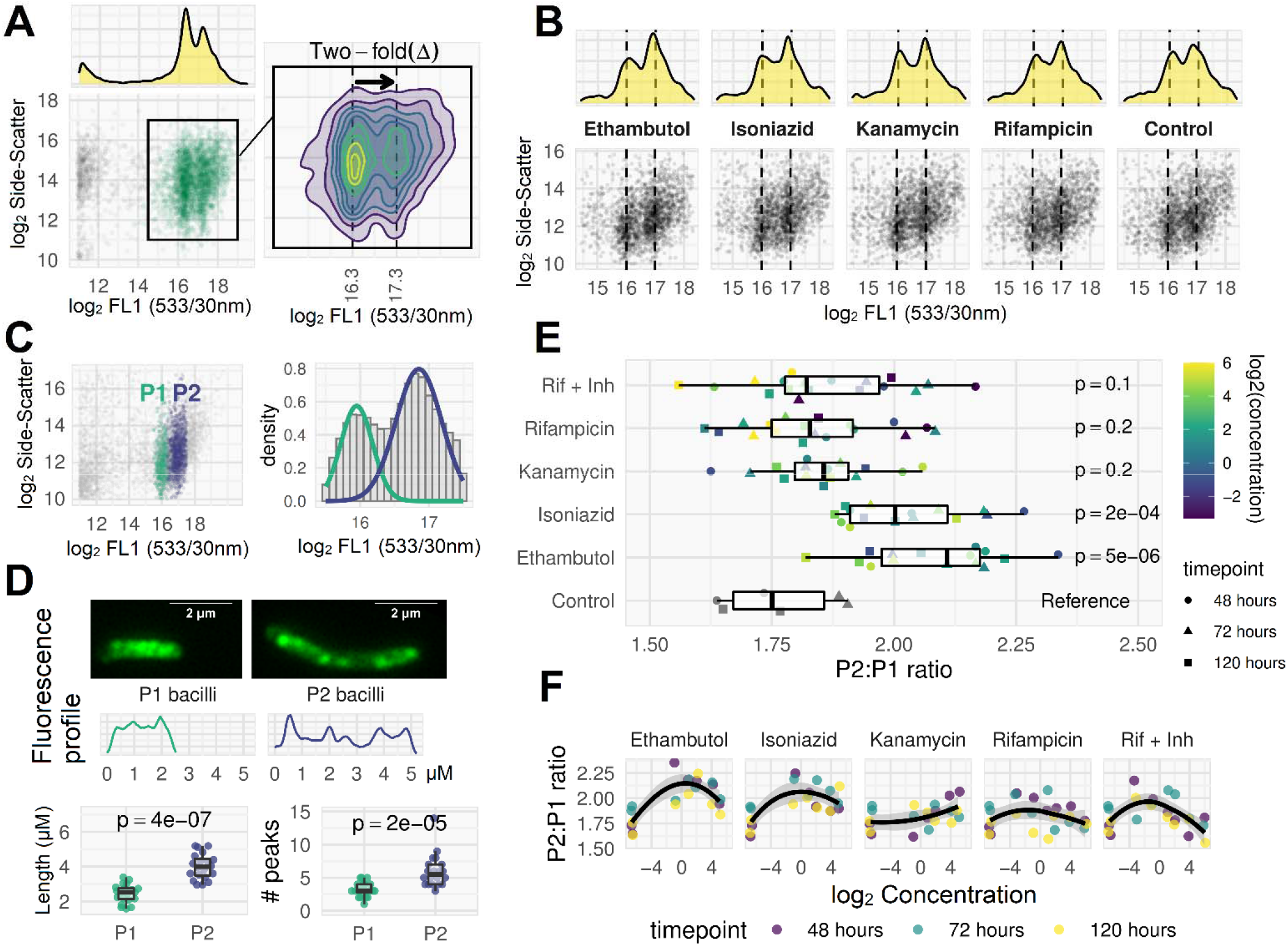
Distinct pharmacodynamics of SYBR-gold-positive sub-populations. **A**. FCM plot for ethambutol-treated (4x MIC, 48 hours), SYBR-gold stained live bacilli. Two discrete sub-populations are visible, separated by approximately two-fold difference in fluorescence. These sub-populations were visible in most ethambutol- or isoniazid-treated cultures, but not readily visible in rifampicin or kanamycin FCM plots (figure 8). **B**. After membrane permeabilization at room temperature, these distinct sub-populations are visible under all conditions including the antimicrobial-free control. **C**. Clustering algorithm (Gaussian mixture model) used to label bacilli as P1 (lower SG fluorescence) or P2 (higher SG fluorescence). **D**. P1 and P2 bacilli sorted for downstream microscopy show different morphologies. A random selection of bacilli images from sorted P1 and P2 sub-populations were measured along their longitudinal axis using ImageJ to assess length and fluorescence profile (two examples shown). P2 bacilli were longer than P1 bacilli (mean 4.Oμm *versus* 2.5μm) and contained approximately double the number of fluorescent ‘peaks’ (mean 6.1 *versus* 3.2; peaks defined by local maxima in a LOESS smoothing function applied to the fluorescence profile plots). **E**. The ratio of P2:P1 bacillary counts was dependent on antimicrobial: isoniazid and ethambutol exposure caused a relative rise in P2 bacilli compared to control, but the same effect was not seen for rifampicin or kanamycin. For the rifampicin plus isoniazid (Rif + Inh) combination treatment, the P2:P1 ratio matched the ratio obtained for rifampicin monotherapy, rather than isoniazid. All p-values were determined from a linear regression of P2:P1 ratio on antimicrobial category, with the antimicrobial-free control as reference category. **F**. Non-parametric Loess regression fitting P2:P1 ratio to log2 concentration (black line with shaded 95% confidence interval) for each antimicrobial condition suggests the pharmacodynamic effect may be non-linearly dependent on concentration.

## Discussion

Detection and accurate quantitation of *Mycobacterium tuberculosis* is fundamental to understanding TB biology. Growing evidence suggests that culture-based methods detect only a sub-population of bacilli,^4,5^ yet these methods remain standard in mycobacterial sciences. By contrast, in response to the analogous problem of differential culturability of microbiota in environmental substrates, FCM has been adopted as an essential method in environmental microbiology research,^60,61^ and industry. ^62,63^ We developed a novel FCM-based method for absolute counting of mycobacteria in liquid cultures. While several groups have reported characterising mycobacteria using FCM, our method is the first to give absolute counts, and can be used to quantify total cell denominator, the presence of cell-clumps, and subpopulations with metabolic activity (using the esterase substrate, Calcein-AM) or membrane permeability (using the nucleic acid stain, SYBR-gold). Our results highlight some critical shortcomings of current ‘gold-standard’ methods for mycobacteria quantification. We also illustrate how the FCM absolute count method can be used for high-throughput, rapid investigations of phenotypic heterogeneity in mycobacteria and demonstrate the capacity to extract pharmacodynamic data using this approach.

Using our FCM method we found that we could reliably identify a subpopulation of the batch culture comprising clumped cells. Further investigations revealed that mycobacterial cultures remain prone to cell clumping in spite of commonly used measures to reduce their formation and to disrupt these before experimentation. We found that sample processing has a major impact on clump-dispersal, such that needle emulsification could increase CFU count approximately two-fold even in early logphase growth. This implies the potential for significant noise and bias in CFU determination, especially for late-log and stationary phase cultures, given that the method depends on serial dilution of dispersed cultures. Our data suggest published protocols^42^ for producing single cell suspensions using centrifugation have no effect on the ratio of clumps to single cells. Using the needle-emulsification and FCM counting methods described would be expected to reduce experimental error in mycobacterial research where bacilli counts are needed to standardise starting conditions, or where the number of bacilli is the dependent variable of interest.

Further, under antimicrobial-free early-mid log-phase growth conditions, CA+ bacillary counts correlate well with CFU counts and can be obtained within 90 minutes using low-cost reagents indicating that the FCM counting method is a practicable alternative to current culture-based methods of estimating cell numbers.

Our FCM method is distinct from previously described mycobacterial FCM protocols primarily because a fluorescence threshold is used to determine when FCM events are recorded. This means that fluorescence-negative events (e.g. a Calcein negative cell) cannot be directly observed but permits accurate absolute counts to be reported for the first time in mycobacterial flow cytometry. Because an absolute cell count denominator can be established with SYBR-gold staining of heat-killed bacilli, the proportion of bacilli with a given characteristic can be ascertained. Importantly, a total cell denominator also allows the proportion of bacilli forming colonies to be measured, which was about 60% in mid-exponential phase of growth in liquid culture.

Our pharmacodynamic results build on previous mycobacterial flow cytometry work reported by Hendon-Dunn *et al.^33^* We replicate their finding that pharmacodynamic flow cytometry profiles based on fluorescent probes of cytoplasmic esterase metabolism and cell wall integrity are different for drugs with different mechanisms of action. We found that the cell wall acting drugs isoniazid and ethambutol were associated with a relatively rapid rise in SG+ cells, while the cytoplasmic targeting rifampicin and kanamycin showed relatively early decline in CA+ bacilli. However, Hendon-Dunn et al. observed only a moderate, concentration-independent effect of rifampicin on Calcein-violet positive bacilli over the first 4 days of exposure. By contrast we found that rifampicin had an early, concentration-dependent effect on CA+ cells, and this effect was substantially greater than for isoniazid. Hendon-Dunn et al. measured relative proportions of Calcein positive and negative cells in cytometry plots, while our method produces absolute cell counts. If antimicrobials have differential effects on the total cell count (which we observed), relative proportions could be unreliable readouts of drug effect (owing to a ‘denominator fallacy’). In addition, the manual gating strategy used by Hendon-Dunn et al. does not appear to capture the shift in mean Calcein fluorescence seen under early rifampicin action, whereas the unsupervised classification approach implemented in this method does.

Again, based on absolute counts, we were able to directly and quantitatively compare CFU and FCM sub-population pharmacodynamics. Under all antimicrobial conditions tested, elimination of colony forming bacilli occurred substantially earlier than the decline in CA+ bacilli or the rise in SG+ bacilli. This means that, at some time points a majority of bacilli are structurally intact with evidence of metabolic activity but do not form colonies. Further, we show that pharmacodynamic effect estimates based on FCM-defined subpopulations give different read-outs from those based on CFU counts: the rifampicin effect on CFU elimination is similar to isoniazid, but rifampicin elimination of CA+ bacilli is markedly greater; rifampicin also has a larger effect on total cell count than the other antimicrobials tested. Rather than simply being a rapid surrogate for CFU counts, the FCM method therefore provides information on antimycobacterial drug pharmacodynamics not captured by CFU counting, but it is unknown if this information is clinically meaningful. Terminally injured bacilli may simply retain metabolic activity with residual enzyme activity in a non-viable cell. Alternatively, as non-growing metabolically active (NGMA) cells can be capable of resuscitation,^55^ this may represent an adaptive response to antimicrobial stress by reducing the physiological consequences of target inhibition.^56^ FCM probes of metabolic and structural integrity would then be more meaningful measures of viability. The latter would be a simple explanation for the lack of correlation between culture-based surrogate endpoints (early bactericidal activity measured using CFU counting, 2-month culture conversion, modelling serial CFU counts or time-to-positivity in liquid culture) and probability of achieving sterilising cure in clinical tuberculosis pharmacodynamics^43^ and warrants testing in clinical samples.

If NGMA bacilli are an adaptive response to antimicrobial exposure then the ability to characterise them using high-throughput methods is critical. By staining live but membrane-permeabilized bacilli with SYBR-gold, we observed two distinct bacilli sub-populations, separated by a two-fold difference in mean fluorescence. We found this phenotype-variation was specifically induced by exposure to isoniazid or ethambutol, but the isoniazid effect was inhibited by the presence of rifampicin. Importantly, the induction of this phenotype could be seen at antimicrobial-condition-timepoints where >99.9% of CFU were already eliminated (e.g. after 72 hours of isoniazid exposure at 4x MIC concentration). After cell-sorting, bacilli from the two-fold brighter subpopulation were found to be longer with double the number of fluorescence foci. Given that SYBR-gold fluoresces when bound to nucleic acid, this implies a bacillary phenotype with double the nucleic acid content, and this phenotypic heterogeneity may therefore represent different numbers of chromosome copies. In a non-human primate model of tuberculosis, “chromosomal equivalents” remain abundant in granulomas that have been sterilised (rendered CFU-negative) by isoniazid therapy.^64^ Peaks separated by a 2-fold difference in fluorescent intensity after staining with ethidium bromide or PicoGreen have been used extensively to define multiple chromosome numbers in *E. coli*^44–46^ Further, several groups have associated polyploidy in *E. coli* with elongated “filamentous” persister cells capable of accelerated antibiotic resistance evolution.^46–48^ In the Wayne model of non-replicating persistence during hypoxia-induced stress, mycobacteria are found to be diploid.^49^ In an *in vitro* foamy-macrophage model, intracellular *Mycobacterium avium* has been shown to enter a reversible dormancy state where the bacilli elongate but do not divide (implying they would not form colonies);^57^ it is suspected that these elongated, metabolically-active but non-replicating mycobacteria may be polyploid.^50^ We speculate that polyploid, metabolically-active but non-colony forming bacilli which are preferentially induced by isoniazid but not rifampicin may be of significant clinical interest. If they represent a drug-tolerant phenotype unobserved by CFU counting, this could explain the fact that, while isoniazid has the most potent early bactericidal activity (EBA, measured by CFU counting), only rifampicin-containing regimens can reliably effect sterilising cure after 6-months (“short-course”) therapy. That drug resistant mutants emerge from phenotypically drug tolerant cells has recently been described for clinical isolates of *Staphylococcus aureus,^58^* a similar mechanism might exist for mycobacteria. The spontaneous drug resistance mutation rates for *M. tuberculosis in vitro* range from 10^−7^ to 10^−9^ and it is somewhat unclear if estimated total mycobacteria numbers *in vivo* allow for development of multidrug resistance through the simple product of these probabilities^59^ – particularly if drug resistance emerges *de novo* after EBA has eliminated most bacilli observable by culture. A population of bacilli, unobserved by CFU counting but capable of elongation and polyploidy, implies ongoing chromosome replication after antimicrobial exposure and a pool of drug-tolerant cells from which drug resistance could emerge.

We used batch cultures in this work which may have limitations. Unexplained variation in FCM sub-population proportions and growth rates between biological replicates, even under antimicrobial-free conditions were seen (e.g. figure 7 B&F). Steady-state cultures – such as the chemostat method used by Hendon-Dunn in their flow-cytometry study – are known to improve reproducibility compared to batch cultures in microbial proteomics and transcriptomics analyses,^51,52^ and are likely to be a major advantage in pharmacodynamic studies. Indeed, cell populations in batch cultures can show complex, non-linear growth patterns in cell size and DNA content^44^ (which are major read-outs from the current implementation of our FCM absolute count method). However these limitations of batch cultures, match those of currently implemented culture methods in research laboratories and are expected to add noise rather than bias to our results.

Overall, our results add to the evidence of limitations in established methods for enumeration of bacilli and support the utility of FCM as a high-throughput, singlecell, culture-independent quantitative tool for the study of mycobacteria in preclinical drug development and ultimately in clinical samples.

## Methods

### Cytometry

Flow cytometry was performed on a BD Accuri™ C6 with manufacturer standard fluorescence detector set-up (FL1, 533/30 nm; FL2, 585/40 nm; FL3, > 670 nm; FL4, 675/25 nm) and data acquisition with BD Accuri™ C6 software including recording processed sample volume. Quality assurance was performed using fluorescent beads as per manufacturer protocol. Manual and extended cleaning cycles were performed at the beginning and end of each flow cytometry session with verification of low event rate in filtered PBS before each run. Cell-sorting experiments were performed on a Bio Rad S3 cell sorter or FACS Vantage with voltage and gain set to recreate BD Accuri C6 plots. All microscopy was performed on a Zeiss Axio Observer 7.

### Sample processing

Needle emusfication was performed with 12 passes through a double luer-lock ended, 25 Gauge, 4-inch, micro-emulsifying needle (CAD7974 Sigma Aldrich). Sonication of cultures prior to FCM to assess effect on clump dispersal was performed by submerging 1ml centrifuge tubes attached to a flotation device in a benchtop ultrasonication water-bath three times for 30 second duration (Ultrawave™ U300HD 30 KHZ; Ultrawave, Cardiff, UK). Heat-kill of mycobacterial samples for “HK counts” was by immersion of aliquots in a waterbath at 60°C for 12 minutes. Removal of antimicrobials prior to CFU plating was by pelleting a 1000μL sample (diluted 2-fold from 500μL with PBS) at 18000g for 12 minutes, removing 900μL of supernatant, resuspending 100μL residual volume in 900μL of 0.22um filtered 0.15% v/v Tween 80 sterile PBS by pipetting, repeated twice (for 10×10 = 100-fold dilution). This will have diluted antimicrobial in solution (unbound) 200-fold prior to plating.

### Culture conditions

Liquid media was prepared from Middlebrook 7H9 media (211887 BD Diagnostics) and 0.22um filtered deionized water according to manufacturer instructions. This was supplemented with 10% v/v Middlebrook OADC (212240 BD Diagnostics), 0.2% v/v glycerol and 0.15% v/v Tween 80. All broth was autoclaved prior to supplementation, and 0.22um filtered prior to use. Liquid cultures were at 37°C in the dark with 150-200rpm agitation in 50ml sterile polyethylene conical flasks in an incubator with an orbital shaking system (model LM-570; MRC Laboratory Instruments Group, London, UK).

Middlebrook 7H10 (262710 BD Diagnostics) agar was prepared with 0.22um filtered deionized water according to manufacturer instructions, with v/v 0.5% glycerol added before autoclave sterilisation. When cooled to 45°C, v/v 10% ADC supplement was added and tri-segmented plates poured to depth 5mm. For CFU counting, 10-fold serial dilutions of samples were prepared in 96-well plates using 0.22um filtered 0.15% v/v Tween 80 sterile PBS. Each segment of a plate was inoculated with 50uL of serial dilutions and spread using disposable, sterile loop spreaders. CFU counts were performed with 3-fold technical replicates and counts averaged. Colony counts between 1 and 100 per segment were accepted and, after adjustment for dilution, averages across dilutions were made where available. Counts were performed on 3 occasions (14, 21, and 28 days) to allow colonies to be counted before overgrowth.

### Reagents

Primary stock dilutions of antibiotic powders were made in 100% DMSO and frozen at-20°C protected from light. Fresh working dilutions were prepared in PBS prior to each experiment, 0.22nm filtered, and stored wrapped in tin foil at 2-5°C refrigeration. Final concentrations of antimicrobials used in *M. bovis* BCG time-kill experiments are reported in multiples of the MIC as indicated. The MIC were: rifampicin, 0.01μg/ml; isoniazid 0.125μg/ml; kanamycin 1.0μg/ml; ethambutol 1.0 μg/ml.

Calcein-AM 50μg vials (ThermoFisher, C3100MP) were reconstituted in 50μL of DMSO on the day of each experiment, further diluted to 200μL with PBS for “working stock”. For cell staining, 5μL of working stock was added per 100μL of sample, with pipette mixing, then incubated in the dark at room temperature for 45-60 minutes before resuspension of bacilli with pipetting. SYBR-gold propriety stock (ThermoFisher, S-11494) was diluted 1000-fold in PBS, aliquoted and frozen at −20°C until use. After thawing aliquot, a further 10-fold dilution (to 10^−4^) in PBS was performed, and 5μL of this working stock added per 100μL of sample to be stained, with pipette mixing, followed by incubation in dark at room temperature for 45-60 minutes before resuspension of bacilli with pipetting.

### Permeabilization of live cells

For the investigation of SG stained subpopulations, published methods for permeabilizing mycobacterial cell walls were reviewed; those with highest reported success and best description of validation^53,54^ were taken forward for testing and adaptation. Permutations of paraformaldehyde / ammonium chloride fixation, ethanol, hydrochloric acid, detergents, and lysozyme were tested at different concentrations, incubation times, and temperatures also assessed iteratively. This led to a final method for reliable permeabilization of BCG bacilli without substantial cell loss, such that over 80% of bacilli could be SYBR-gold stained (compared to the heat-killed total cell count denominator gold standard). In the final method, a 500μL sample was diluted to 1ml with PBS v/v 0.15%Tween 80, without wash step or fixation. After needle emulsification, lysozyme was added to final conc 0.1mg/ml, and the sample incubated for 45 minutes at 37°C. 500μl triton-X-100 was then added to final concentration v/v 0.2%. This was pelleted (16000*g*, 5 min) and re-suspended in 500μl PBS-tween; 40μl working stock SYBR-gold added and incubated at room temperature for 2-4 hours.

### Data analysis

Raw flow-cytometry data was extracted from .fcs files exported from BD Accuri C6 software using *flowCore* (v 2.0.1) Bioconductor package^65^ and all analysis performed in Rstudio v1.1.463. Rather than using manually placed gates to classify events, unsupervised machine-learning classification algorithms were used. For main flow cytometry plots (figure 8) k-means clustering was applied to FL1 height, FL1 area, and Side-Scatter height observations from one replicate using kmeans() function in *stats* package^66^ in R, and the clustering solution applied to all the data. Optimal number of clusters was determined empirically using NbClust() function from *NBClust* package^67^ in R. To separate P2 and P1 events in permeabilized SG-stained live cells (figure 11), a Gaussian Mixture Model was fit to all the data with 2 component distributions using normalmixEM() function in *mixtools* package.^68^ In all FCM scatter plots, log transformations to base e (natural logarithms) are presented unless otherwise indicated (plots with log transformations to base 2 are used in cases where a doubling of fluorescence is a specific feature of interest).

Time-kill curves were summarised in descriptive plots (figure 9) using non-parametric loess regression models. To extract summary measures of antimicrobial effect, the time-kill data was modelled using a linear mixed-effects model, with a fixed effect of intercept, and random slopes for antibiotic condition and replicate:

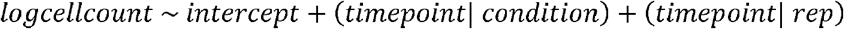

This captures the crossed experimental design where each replicate was assessed under each antimicrobial condition, and each replicate assessed under each condition. The antibiotic condition effect, defined as slope gradient for the time-kill curve, was then extracted, independent of replicate effect, from the model as a summary PD measure. Because the dependent variable is on a log scale this assumes a mono-exponential decline in cell populations under antimicrobial action. The dependent variables assessed were CFU count, CA+ count, HK count, and proportion SG+. The R package *lme4* was used for this modelling.^69^

Antimicrobial effects extracted from these models were related to drug concentration for each antimicrobial using a standard sigmoid E_max_ PK/PD model, of form:

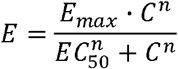

Where *E* is the PD effect (the slope gradient estimates by mixed-effects modelling above), *C* is the drug concentration (known from experimental condition), and the remaining parameters are estimated from the data: *E_max_* (maximum achievable effect of antimicrobial), *EC_50_* (the drug concentration where half of *Emax* is obtained), and *n* (a scaling parameter). Models were fitted using non-linear least squares (nls() function in R.

Fluorescence profiles of bacilli (figure 11D) were extracted as .csv files from microscopy images using Fiji (ImageJ).^70^ This raw data was processed using a custom-built function defining local maxima in smoothed profiles to count fluorescent peaks.

## Acknowledgments

The work was supported by Wellcome Trust fellowship 105165/Z/14/A.

RJW receives support from Francis Crick Institute which is funded by Wellcome Trust [FL0010218], UKRI [FC0010218], CRUK [FC0010218] and also support from Wellcome Trust [104803, 203135].

## Author contributions

DAB, GD, DL, RJW, and GM conceived of the approach and initiated the project. DAB, DFW, VM, GD and GM conceived specific methods and experiments. DAB, CO, AK, and MKM designed and performed the experiments. DAB analysed data and wrote the manuscript with input from all authors.

## Data availability

Data and analysis scripts are available in an Open Science Framework repository here https://osf.io/gwhpd/?view_only=02a61e8134cc4adca3840df604e0e38b

